# The subcellular architecture of the *xyl* gene expression flow of the TOL catabolic plasmid of *Pseudomonas putida* mt-2

**DOI:** 10.1101/2020.08.30.273938

**Authors:** Juhyun Kim, Angel Goñi-Moreno, Víctor de Lorenzo

**Author notes:** Correspondence to: Víctor de Lorenzo, Centro Nacional de Biotecnología-CSIC, Campus de Cantoblanco, Madrid 28049, Spain. Department of Chemical Engineering, Imperial College London, London, SW7 2AZ, UK.

## Abstract

Despite intensive research on the biochemical and regulatory features of the archetypal catabolic TOL system borne by pWW0 of *Pseudomonas putida* mt-2, the physical arrangement and tridimensional logic of the *xyl* gene expression flow remains unknown. In this work, the spatial distribution of specific *xyl* mRNAs with respect to the host nucleoid, the TOL plasmid and the ribosomal pool has been investigated. *In situ* hybridization of target transcripts with fluorescent oligonucleotide probes revealed that *xyl* mRNAs cluster in discrete foci, adjacent but clearly separated from the TOL plasmid and the cell nucleoid. Also, they co-localize with ribosome-rich domains of the intracellular milieu. This arrangement was kept even when the *xyl* genes were artificially relocated at different chromosomal locations. The same happened when genes were expressed through a heterologous T7 polymerase-based system, which originated mRNA foci outside the DNA. In contrast, rifampicin treatment, known to ease crowding, blurred the confinement of *xyl* transcripts. This suggested that *xyl* mRNAs intrinsically run away from their initiation sites to ribosome-rich points for translation—rather than being translated coupled to transcription. Moreover, the results suggest that the distinct subcellular motion of *xyl* mRNAs results both from innate properties of the sequence at stake and the physical forces that keep the ribosomal pool away from the nucleoid in *P. putida*. This scenario is discussed on the background of current knowledge on the 3D organization of the gene expression flow in other bacteria and the environmental lifestyle of this soil microorganism.

**IMPORTANCE:** The transfer of information between DNA, RNA and proteins in a bacterium is often compared to the decoding of a piece of software in a computer. However, the tridimensional layout and the relational logic of the cognate biological hardware i.e. the nucleoid, the RNA polymerase and the ribosomes, are habitually taken for granted. In this work we inspected the localization and fate of the transcripts that stem from the archetypal biodegradative plasmid pWW0 of soil bacterium *Pseudomonas putida* KT2440 through the non-homogenous milieu of the bacterial cytoplasm. The results expose that— similarly to computers also—the material components that enable the expression flow are well separated physically and they decipher the sequences through a distinct tridimensional arrangement with no indication of transcription/translation coupling. We argue that the resulting subcellular architecture enters an extra regulatory layer that obeys a species-specific positional code that accompanies the environmental lifestyle of this bacterium.

## INTRODUCTION

The TOL system encoded by plasmid pWW0 of *Pseudomonas putida* mt-2 is to this date the most thoroughly studied example of biodegradative system in soil microorganisms. The primary function of this catabolic device is enabling carrier bacteria to grow on toluene, *m-*xylene, *p-*xylene and other related aromatics through a set of enzymes encoded by *upper* and *lower* plasmid-borne operons (Fig. 1A; 1-3). While catabolic traits of this sort are not uncommon in many other environmental isolates, what makes the TOL system special is the extraordinary regulatory intricacy that controls expression of the *xyl* genes and their high-level interplay with the host’s physiological regulons. The many mechanisms unveiled over the years in this respect seem to include nearly every device known in the prokaryotic word for ruling the gene expression flow (4-6). This state of affairs has made the TOL plasmid and its bearer (*P. putida* KT2440) a beneficiary of the suite of conceptual and material tools of contemporary Systems and Synthetic Biology (7, 8). In particular, the wealth of experimental data on expression of the *xyl* genes has enabled the understanding of the cognate regulatory network as a complex device that processes inputs into outputs following a layer of logic gates implemented with promoters, transcriptional factors and sRNAs (9, 10). It could thus be argued that we know at this point much about the genetically-encoded *software* of the system and the relational logic that rules its performance. In contrast, we know virtually nothing of the physical arrangement of the *hardware* that sustains the same process. In particular, the gross spatial disposition of the molecular actors that execute the transfer of information from the *xyl* genes to production of catabolic enzymes is unknown.

**FIG 1.**
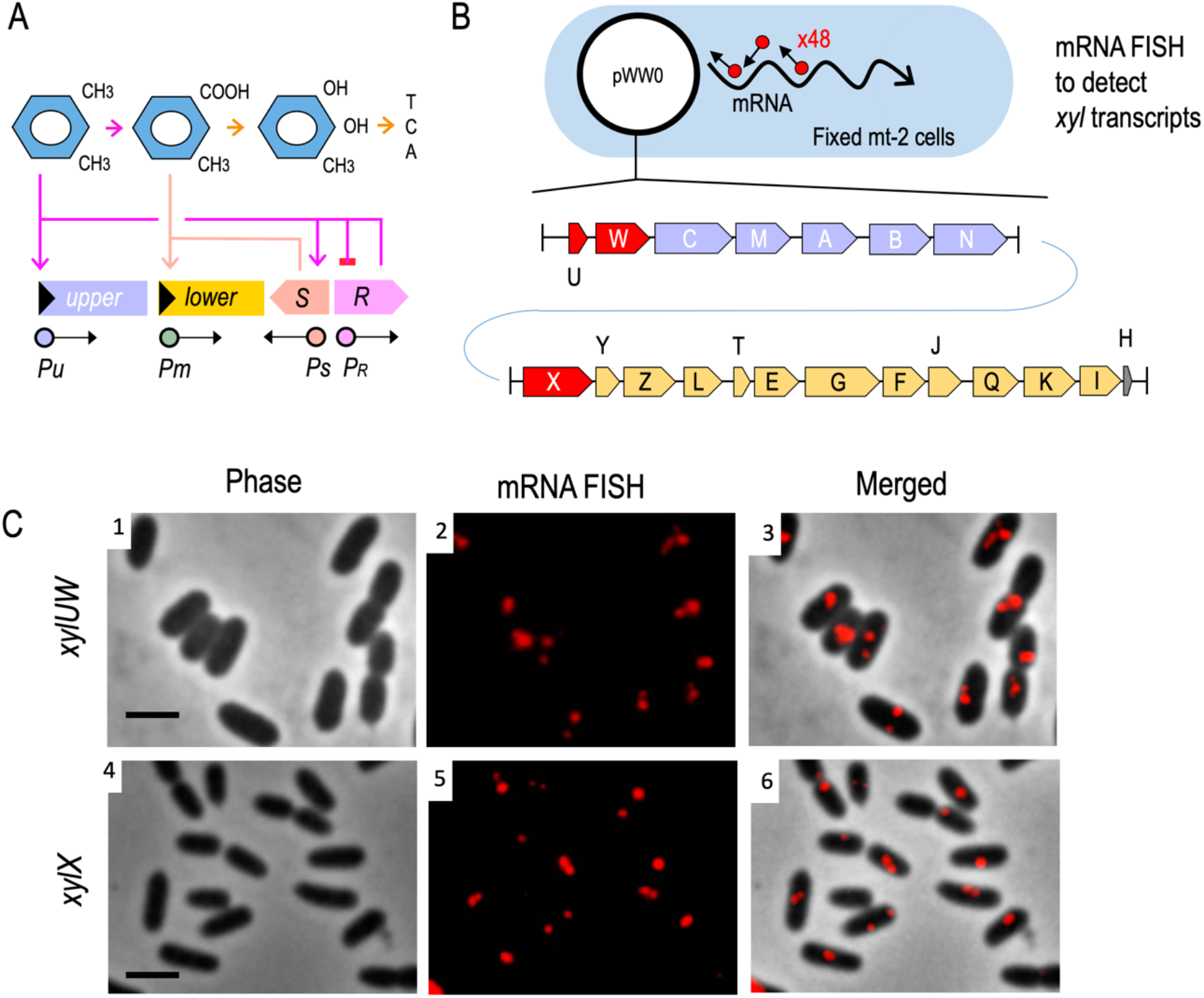
Interplay between metabolic and regulatory components of the TOL network and visualization of the transcripts from either the *upper* or the *lower* of the system. (A) In the presence of *m*-xylene, XylR (expressed from P_R_) activates both promoters *Pu* (which transcribes the *upper* pathway operon) and *Ps*, stimulating expression of *xylS* gene. Also, XylR expression is negatively auto-regulated. In turn, the XylS protein induces the lower operon by activating *Pm* promoter with its effector 3MBz as the metabolite of *m*-xylene. (B) Scheme showing the fluorescent *in situ* hybridization (FISH) experiment to visualize mRNAs from the TOL operons of plasmid pWW0. For this experiment, *P. putida* mt-2 strain was grown in succinate-supplemented minimal medium to the exponential phase, then further cultured for 2 h with saturating vapors of *m*-xylene to activate the *xyl* genes. After the fixation of the sample with formaldehyde, the FISH experiment was performed with either the *upper* (*xylUW*) or the *lower* pathway (*xylX)* probe set, involving the red fluorophore-labeled 48 oligos complementary to each mRNA sequence. (C) Microscope images of the FISH approach responding to the *xylUW* mRNA and the *xylX* mRNA, respectively. Phase-contrast (panel 1 and 4), mRNA-red signals (panel 2 and 5), and the composite images (panel 3 and 6) are shown. Scale bar, 2.5 µm.

The same sort of questions has been tackled in other prokaryotic systems. There seem to be at least two different patterns of transcript localization. In one scenario, the mRNA remains close to the site of transcription. This is the case of the mRNAs of *groESL* and *creS* of *Caulobacter crescentus* and *lacZ* of *E. coli*, which appear to remain in the vicinity of the corresponding genomic DNA loci where they are transcribed (11). This suggests that for mRNAs to be translated they need to be associated to the ribosomes while being produced (as the prevailing view of generalized transcription-translation coupling would imply) or short after their creation by RNAP (11, 12). An alternative scenario involves migration of mRNAs to specific spots of the cytoplasm for translation close to the site(s) where the gene products are needed. While this setting argues against generalized transcription/coupling translation, there is solid evidence of its occurrence e.g. the *bglGFB* operon, whose products were located in sub-cellular localizations where cognate proteins were expected to work (13). This example may not be anecdotal as transcriptome-wide scale studies of mRNA localization in *E. coli* revealed that a large share of transcripts were found in cell domains (e.g. membranes, cytoplasm, poles) that coincided with the function of the cognate proteome (14, 15). Furthermore, the mRNA of archetypal membrane proteins LacY and TetA are distinctively close to the cell envelope (16, 17). In other instances, a signal-recognition particle (SRP) is involved in the movement of mRNAs to the membrane, as translation of some mRNAs encoding inner-membrane proteins produces a signal peptide that recruits such SRP and the complex leads the transcript to its intracellular address (15, 16, 18). Finally, Rho-dependent transcription termination in *Bacillus subtilis* is somewhat weak, and *runaway*, untranslated mRNAs are abundant (19). In sum, the fate of each transcript seems to be both gene (i.e. sequence)-dependent and species-dependent.

In this work we have inspected the localization of mRNAs initiated in the catabolic promoters of the TOL plasmid pWW0 of *P. putida* mt-2. The starting point for tackling the issue is the earlier observation that RNA polymerase (RNAP) of *P. putida* fully co-localizes with the chromosomal DNA of the nucleoid while being entirely apart from the bulk of the ribosomal pool (20). This observation suggested that rather than being coupled to translation *ab initio*, many (if not most) of the mRNAs initiated at chromosomal promoters need to move to ribosome-rich, nucleoid-free domains of the intracellular space for translation. Note that in the case of the *xyl* promoters, transcription initiates in an extrachromosomal element, and therefore the 3D itinerary of the corresponding mRNAs could be different. As shown below, by merging genetic analyses with *in situ* RNA-FISH and DNA-FISH technology we could faithfully locate the relative positions of the nucleoid DNA, the pWW0 plasmid, the *xyl* transcripts and the ribosomes and predict the motion of cognate mRNAs through the cell interior. The results exposed an unexpected degree of physical partition among the material actuators of the gene expression flow that may help *P. putida* to deal with its typical environmental settings.

## RESULTS AND DISCUSSION

### *mRNAs of catabolic genes expressed from* the *TOL plasmid are spatially organized*

To visualize specific mRNAs of *P. putida* mt-2 stemming from the TOL operons of plasmid pWW0 we adopted an RNA FISH approach (Fig. 1B). To this end, the strain was cultured in M9 minimal medium with succinate as the sole carbon source until the cells reached exponential phase. At this point the cultures were exposed (or not) to saturating vapors of *m*-xylene to induce transcription of the catabolic *xyl* genes. After 2 h, samples were collected and fixed with formaldehyde for hybridization with specific probes as described in the Materials and Methods section. For this, two sets of 48 CAL Fluor Red 610-tagged fluorescent oligonucleotides (20 nt long; Supplementary Table S2) were synthesized that covered, respectively, the leading 1418 bps of the *upper* TOL operon transcript spanning the whole of *xylU* and part of *xylW* (encoding benzyl alcohol dehydrogenase) and the front segment of the TOL *lower* operon, encompassing 1203 pb of the *xylX* gene (alpha subunit of toluate 1,2-dioxygenase). After hybridization with these oligo sets, the samples were washed and red signals inspected with fluorescence microscopy.

As shown in Fig. 1C, distinct, discrete fluorescent foci were clearly noticed under the microscope (1-2 per cell) from the cultures subject to *m*-xylene exposure upon in situ hybridization with *upper*-pathway or *lower*-pathway specific oligonucleotides. In contrast, no signals were detected in bacteria grown in M9-succinate without aromatic effector (Supplementary Fig. S1A). These observations accredited that the designed probe sets and the methodology proper were working as expected, as there were no signals that could be attributed to hybridization with DNA. Therefore, the foci appeared to represent *upper* or *lower xyl* transcripts. To further benchmark the experimental approach, cells were treated with either toluene (an alternative TOL substrate) or *o-*xylene (a gratuitous inducer of the *upper* pathway and downstream activation of the *lower* pathway (21, 22; Fig. 1A). Under these conditions, fluorescent foci for both the *xylUW* and the *xylX* transcripts were expectedly observed. In contrast, cells treated with benzoate or 3-methyl benzoate (substrates of the *lower* pathway) produced signals for *xylX* mRNA only (Supplementary Fig. S1B). Appearance of the fluorescent signals matched the known regulatory network that rules the interplay between the substrates, regulators and catabolic operons of the TOL system (Fig. 1A). We could thus safely consider that the foci inside cells shown in Fig. 1C were the result of the *bona fide* hybridization of the fluorescent probes to specific mRNA of *xylUW* or *xylX*. Closer inspection of the images revealed that no red output ever appeared dispersed throughout the cell, but always as discrete foci (Fig. 1B). Yet, their position in respect to the cell shape varied among cells and signals were located near the center, the poles, the contour or the septum of the cells (Fig. 1C). We thus set out to characterize this asymmetrical localization of the *xyl* transcripts in respect to the nucleoid occupation and to the TOL plasmid, as explained below.

### *xyl* mRNAs occupy subcellular nucleoid-free regions

In order to identify the relative localization of the *xyl* transcripts in respect to the nucleoid, following hybridization with the fluorescent probes described above the bacterial genome was stained with 4’, 6-diamidino-2-phenylindole (DAPI) in cells exposed to *m-*xylene. The results of this procedure with the *upper* and *lower* pathway TOL probes (*xylUW* and *xylX*) respectively are shown in Fig. 2. Images were processed and signals analyzed with the CellShape software. For interpreting these results, note that [i] previous work showed that the DNA of the nucleoid overlaps spatially with *P. putida*’s RNAP (20) and [ii] DAPI binds both chromosomal and plasmid DNA. Inspection of the cells under the microscope (Fig. 2A) revealed that signals stemming from *xylUW* or *xylX* transcripts (red foci) were located in sites inside cells with a low DAPI signal, i.e. in the nucleoid-free regions. In order to quantity the phenomenon, >100 pictures of individual DAPI-stained and fluorescent oligo-hybridized cells were separately recorded with the blue and red channels of the fluorescence microscope and they were automatically inspected with the CellShape image analysis tool (23; Fig. 2B). The outcome of this analysis is shown in Fig. 2C. Very few red foci (< 1%) overlapped with the densest DAPI signals and as little as 15% of fluorescent spots from either *xylUW* or *xylX* RNA were detected at the peripheral regions of genomic DNA. Instead, the vast majority of the remaining foci were placed away from the blue signal (Fig. 2C). It thus looked like the bulk of TOL transcripts were located within subcellular regions with no or little overlap with the DNA signal and therefore virtually devoid of RNAP (20). This suggests that once formed, *xyl* mRNAs could migrate for translation to a site different from the place where transcription is initiated. Note that the pathway substrate (*m-*xylene) dissolves in the cell membrane (24). Moreover, the *xylM* product (hydroxylase component of the leading pathway enzyme xylene monooxygenase) is located in the membrane (25). It is not uncommon that genomic sites encoding envelope-associated proteins are transcribed coupled and co-translationally inserted to the membrane through a so-called *transertion* mechanism (16). In the case of the TOL transcripts it looked instead that movement away from the DNA begins after transcription has terminated and the mRNA has disengaged from the nucleoid. To clarify this, we examined the relative position of the *xyl* mRNA in respect to their actual origin inside the cells (i.e. the TOL plasmid) as explained below.

**FIG 2.**
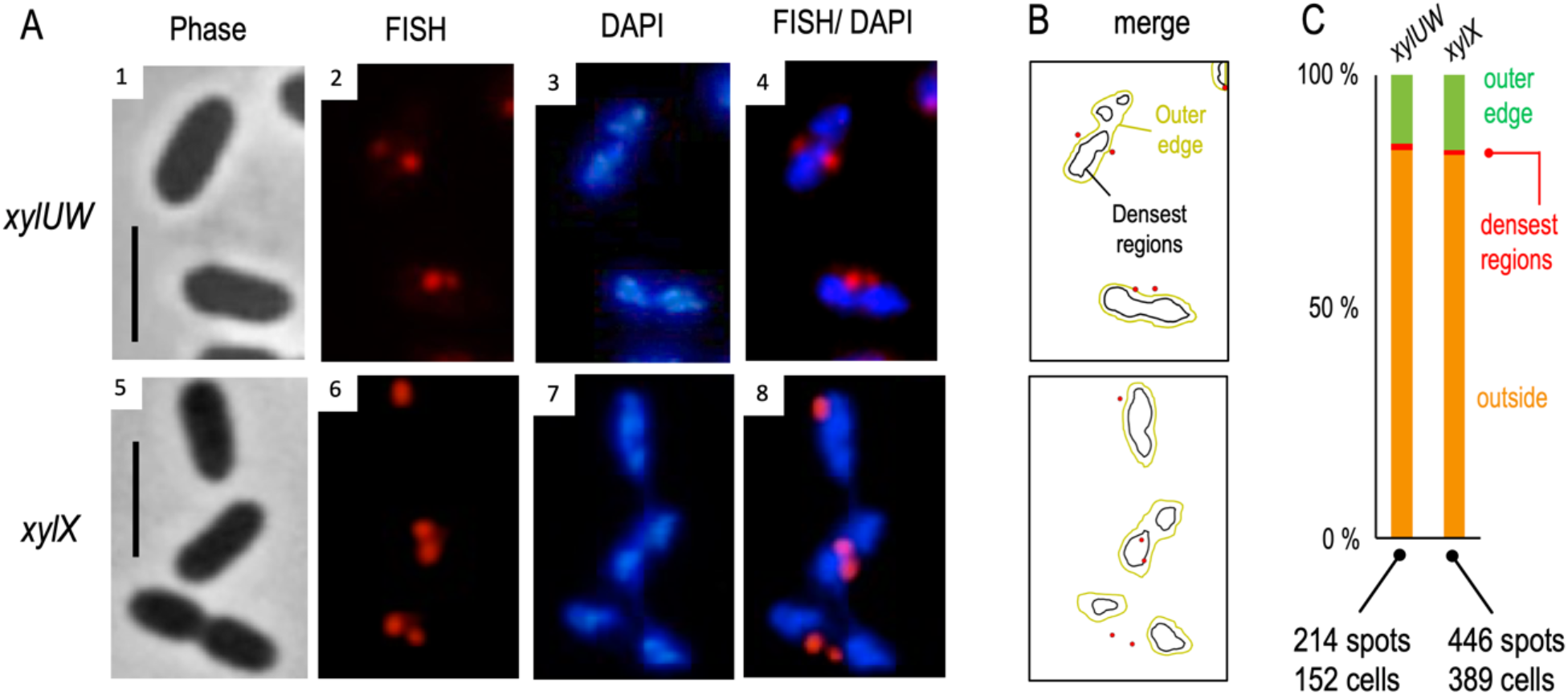
Spatial distribution of the TOL catabolic transcripts with respect to the nucleoid occupation within cells. (A) The mRNA signals, against the *xylUW* or the *xylX*, were obtained by the FISH assay (panel 2 and 6), and the nucleoid was stained with DAPI (panel 3 and 7) in the procedure of the experiment to examine distribution of DNA. The mRNA-red and the DNA-blue channels were merged (panel 4 and 8) to identify relative position of the transcript with respect to the nucleoid. Phase-contrast images (panel 1 and 5) corresponding to the red and the blue channels are also shown. Scale bar, 2.5 µm. (B) Using the image analysis tool as CellShape, fluorescent signals such as mRNA red spots and the nucleoid contour were identified from exemplary cells appeared in (A). The densest regions of the DAPI-stained regions are within black lines, while relatively less concentrated DNA regions are depicted with yellow lines. (C) The abundance of *xyl* mRNAs in the nucleoid-free regions. The *xylUW* (214 spots from 152 cells) or the *xylX* signals (446 spots from 389 cells) were analyzed concerning the distribution of the nucleoid.

### xyl transcripts map near but do not overlain the TOL plasmid

Subcellular localization of the TOL plasmid within the whole of the DAPI-stained DNA of the *P. putida*’s nucleoid required first tagging pWW0 with a different fluorescent label. A 9679 pb DNA sequence including an array of tandemly repeated *tetO* operators for the TetR repressor (26) was inserted in a site of the plasmid between and close to the *upper* and *lower* operons i.e. by ORF105 of the current annotation (27); Supplementary Fig. S2A). The corresponding sequences could then be exposed through FISH with a special type of 6-Carboxyfluorescein (6-FAM)-labeled 18 nt oligonucleotides. These encoded the *tetO* operator and bore a distinctive chemical configuration (so called locked nucleic acid LNA structure) that increases specificity for target DNA (11; Supplementary Fig. 2A). Samples were thus first hybridized with *xyl*-specific (red fluorescence) and pWW0-specific (green fluorescence) probes and then stained with DAPI. As a result of this procedure, both red, green and blue fluorescent signals could be mapped in cells exposed to *m*-xylene, corresponding to *xylUW* or *xylX* mRNA, plasmid, and nucleoid, respectively (Fig. 3B-D and Fig. 4B-D). In contrast, cells grown without the aromatic compound lacked the red signal altogether (Supplementary Fig. S3A).

**FIG 3.**
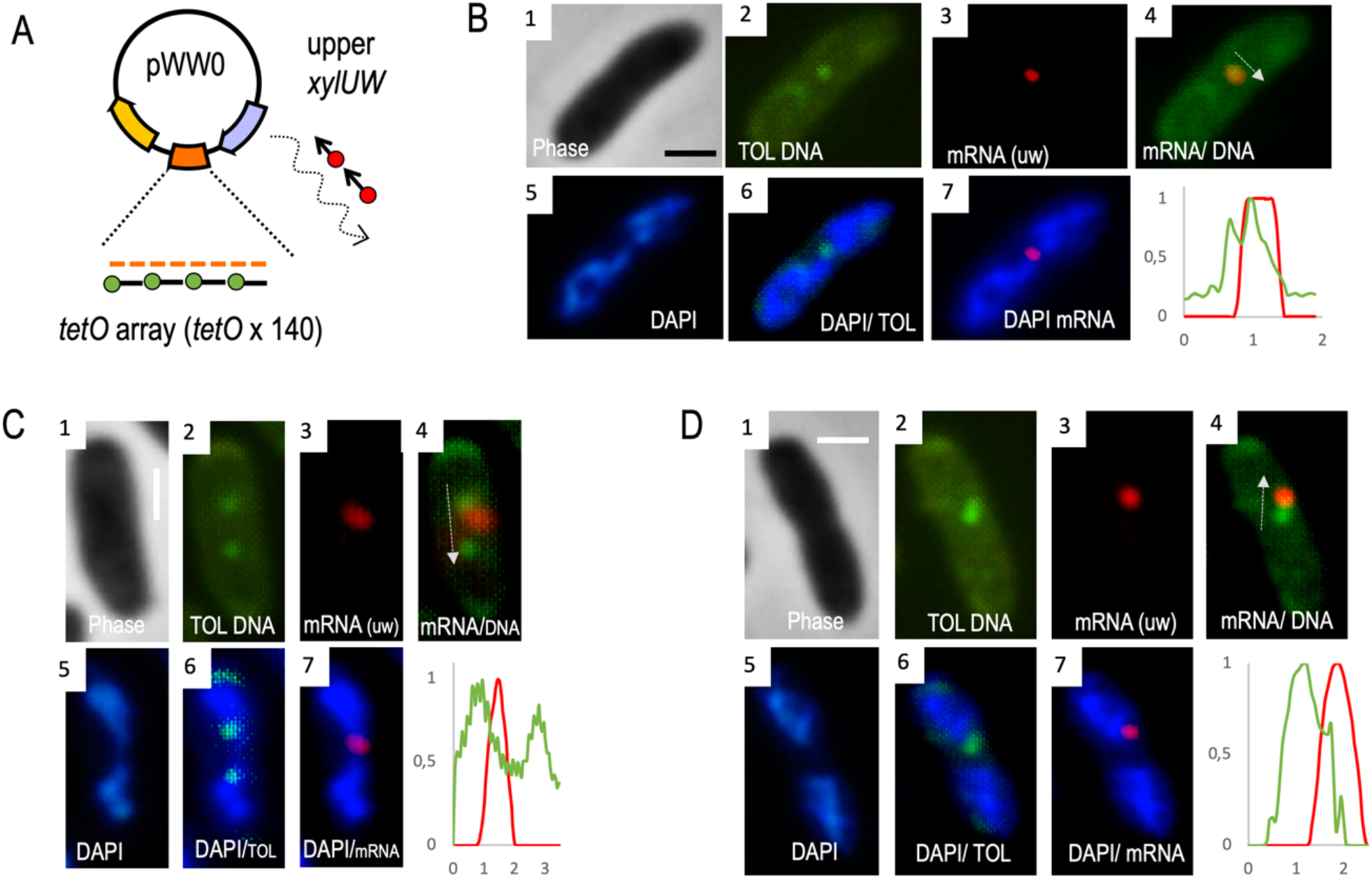
Co-visualization of the *xylUW* transcript and the pWW0 plasmid. (A) Sequentially combined RNA-DNA FISH was carried out to visualize both the *xylUW* mRNA and the pWW0 plasmid in the cell. In order to achieve DNA signals with the approach, the tandem copies of the *tet operators* were inserted into the *orf105* locus between the *upper* and *lower* operons and the green-fluorophore tagged *tetO* probe was used for the hybridization in DNA-FISH procedure. (B-D) The mt-2 (pTOL-tetO) strain, exposed to *m-*xylene for 2 h, was fixed for the FISH approach. The resulting images from the experiment were shown with representative cells demonstrating dynamic localization patterns of the mRNA relative to the DNA locus. The microscopy displays phase-contrast (panel 1), the plasmid DNA (green; panel 2), the *xylUW* mRNA (red; panel 3) and the nucleoid (blue; panel 5), respectively. The composite images such as the mRNA/ the plasmid DNA (panel 4), the plasmid DNA/ the nucleoid (panel 6), and the mRNA/ the nucleoid (panel 7) are also shown. Relative intensity was measured along with the mRNA-red and the plasmid-green signals: y-axis refers to relative fluorescent intensity; x-axis indicates pixel distance representing the arrow drawn on the composite image (panel 4). Scale bar, 1 µm. As a sidelight, the images of panels #2 show also that the TOL plasmid exists as a low-copy replicon (∼ 1 molecule) per *cell* of *P. putida*.

**FIG 4.**
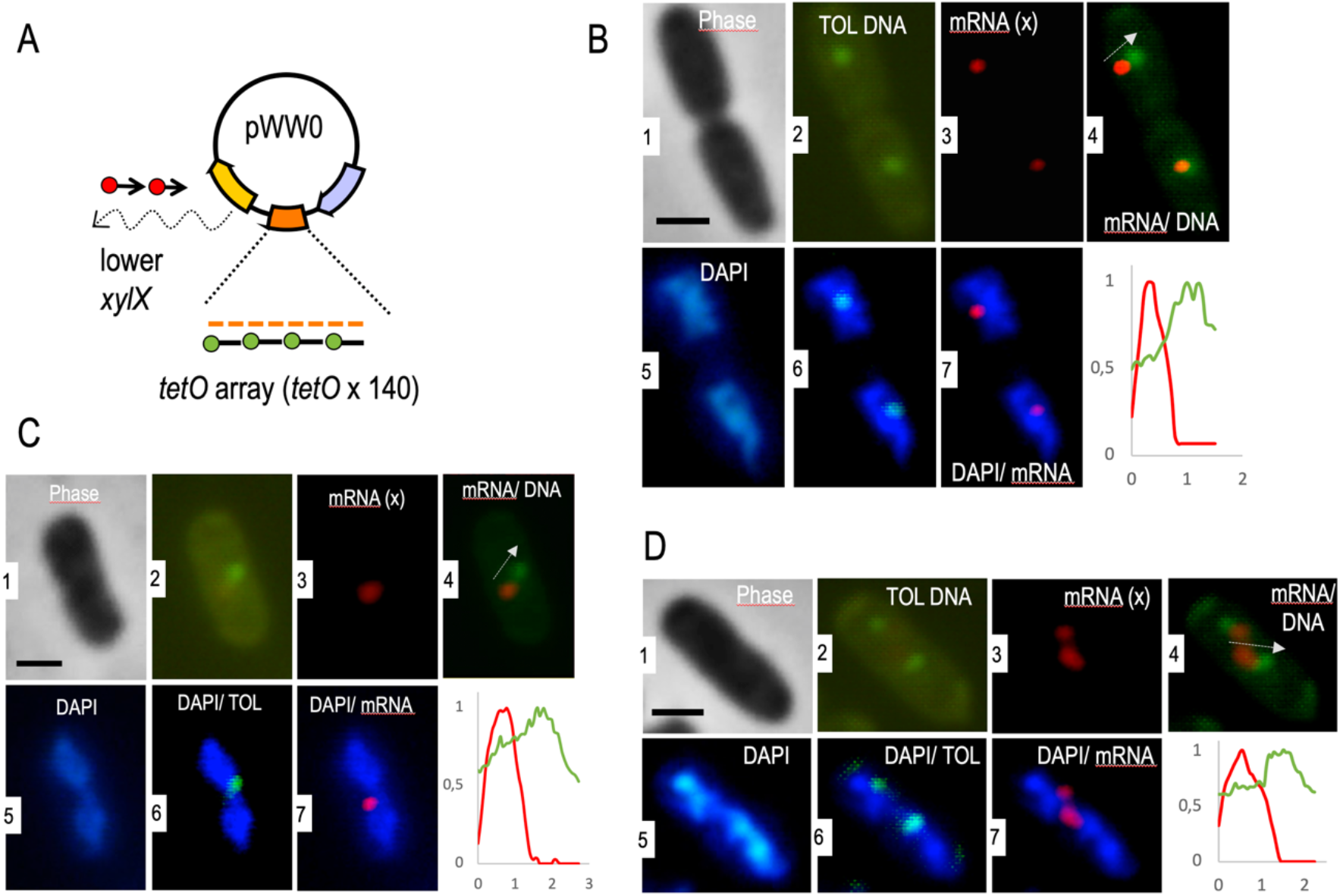
Dual labeling of the *xylX* mRNA and the pWW0 plasmid. (A) The same approach as described in Figure 3A, but red fluorophore-tagged *xylX* probes were applied for the combined RNA-DNA FISH to visualize both the *lower pathway* transcripts and the TOL plasmid. (B-D) Cellular position of the plasmid DNA and the *xylX* mRNA was detected in the example cells by employing the FISH approach. All panels are organized as the same statement of Figure 3.

The control images (panels 1 and 2 of Figs. 3B-D and 4B-D; Supplementary Fig. S3) show individual bacteria hosting discrete green spots in cells bearing the TOL plasmid variant tagged with *tetO* array. In contrast, no defined green foci were observed in cells with the intact TOL plasmid devoid of the same array (Supplementary Fig. S2B). This verified that the *tetO* probe hybridized specifically to the tagged plasmid without significantly interact with any other DNA of the cells. Further inspection of the pictures indicated that the green signals appeared only once or twice in every cell (in the last case in a symmetrical fashion; Supplementary Fig. S3). Such a subcellular distribution of the plasmid has been observed before (28) and plausibly reflects the plasmid partition system (29-31). Once the position of the TOL plasmid was determined, by filtering the images with the adequate color channels we could pinpoint the relative localization of pWW0, the nucleoid and the TOL transcripts.

Fig. 3B-D and Fig. 4B-D show a few examples of thereby processed cells to locate the *xylUW* (*upper* pathway) and the *xylX* mRNAs (*lower* pathway), respectively. Each of these images was not only visually inspected but also the signals quantified with the CellShape software (23; Supplementary Fig. S4). In general, the transcripts from either the *upper* or the *lower* TOL pathway behaved similarly. The first piece of information resulting from a detailed image analysis (panels 2/6 of Fig. 3B-D and Fig. 4B-D) is that TOL plasmid could be found both in DAPI-intense regions and in the peripheral space of the cytoplasm with little or insignificant DAPI signal. This agrees with the pWW0 predicted segregation system: the presence of plasmid-encoded *parA* and *parB* indicates pWW0 to bear a Type I partition mechanism (27) though which the plasmid can either detach or remain connected to the chromosome through a ParA-ATP-nucleoid complex (30).

Inspection of colored spots merged within the contour of single cells exposed two predominant spatial arrangements of the different signals. In one case, red and green spots virtually co-localized (e.g. Fig. 3B), while in most others, the two signals were close to each other but clearly separated (Fig. 3C-D and Fig. 4B-D). It is likely that these two scenarios reflect different stages of transcription of *xyl* genes. In an early stage, mRNA is necessarily tethered to its DNA template in the plasmid and therefore the gene and its transcript occupy the same spot. But at a later stage the *xyl* mRNA could move away from the plasmid towards nucleoid-free ribosome-rich domains of the cell inside (20). Note that the sizes of the *upper*- and *lower* TOL operons are ∼ 8 Kb and 11 Kb respectively and, if linearly stretched, the 5’ ends of their full-length mRNAs could be found well away from the plasmid while still tethered to the template. However, it is known that the mRNA from the *meta* pathway is quite short lived and starts being degraded before it is fully synthesized (32). Note also that mRNA tends to form secondary structures that shorten the distances between the 5′ and 3′ ends (33). On this basis, we argue that separation of red and green spots in the images shown in Fig. 3, Fig. 4 and Supplementary Fig. S4 reflect an authentic migration of the TOL transcripts away from their DNA template. The next obvious question was whether this was result of TOL genes being encoded in a type of plasmid that largely stays in the periphery of the nucleoid (see above). Alternatively, the unexpected localization of *xyl* mRNAs could stem from intrinsic properties of the transcripts themselves. In order to sample these possibilities we entered various types of perturbations in the system as described below.

### xyl transcripts move away from the nucleoid regardless of their replicon

Although the native physical location of the TOL genes is in plasmid pWW0, the same growth phenotype on aromatic compounds can be brought about when the *upper* and the *lower* pathways are placed in the chromosome. This may happen either naturally (34) or by engineering the corresponding DNA segments in the genome of a heterologous host (35). We took advantage of this for exploring whether the position of the *xyl* DNA template affects the spatial distribution of the thereby transcribed RNAs. To this end, we used *P. putida* PaW140 (36), which carries both the *upper* and *lower* operons and their cognate regulators in its chromosome (Fig. 5A). The strain was grown and induced with *m*-xylene under identical conditions as before and cells fixed and hybridized with RNA probes to expose *xylUW* and *xylX* transcripts in respect to the bacterial nucleoid. The results of Fig. 5B showed the number of foci per cell to increase by ∼ 30% in *P. putida* Paw140 as compared to the lower numbers of the pWW0-bearing *P. putida* mt-2 strain (Fig. 2C), surely due to a higher transcriptional activity. This is not surprising as the pedigree of strain *P. putida* Paw140 involved random chromosomal insertion of TOL genes and selection for best growers on *m-* xylene, what plausibly favored implantation in regions of high transcriptional activity (37-39). Yet, when RNA-red signals were compared to those of the DAPI-stained nucleoid, most of them were observed away of the denser DNA regions. Specifically, < 3% of the red spots of either *xyl* mRNAs overlapped the more compacted chromosome, while 74% of *xylUW* and 87% of *xylX* signals were enriched in the peripheral space of the cell (Fig. 5B and 5C). Preferential localization of *xyl* mRNAs was thus maintained regardless of the replicon (plasmid of chromosome) that bears the cognate genes. Moreover, the higher incidence of TOL mRNAs in the nucleoid-free subcellular regions corroborates that *xyl* transcripts move away from their transcription site.

**FIG 5.**
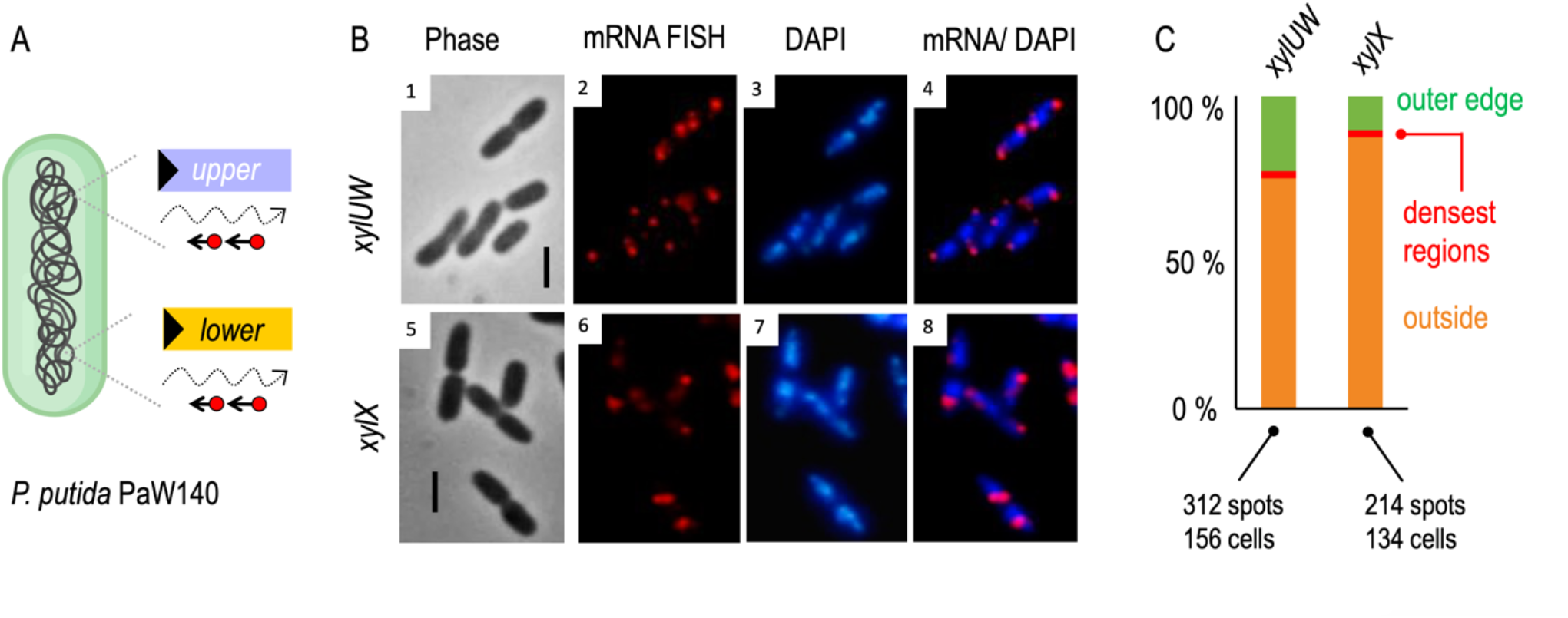
Effect of the replicon type on the localization of *xyl* mRNA. (A) The Paw140 strain, carrying the entire TOL system in the chromosome, was used to identify the distribution of *xyl* mRNAs by following the same procedure applied for the mt-2 strain. (B) RNA-FISH microscopy enabled to visualize mRNA signals corresponding to the *xylUW* (panel 2) and *xylX* mRNA (panel 6), respectively. Their counterpart images such as phase contrast (panel 1 and 5), the nucleoid (blue; panel 3 and 6), and composite images (panel 4 and 8) are shown. Scale bar, 2.5 µm. (C) Quantification of mRNA-red foci with respect to the nucleoid occupation. Note that the number of red stop per cell increased by ∼ 30% in *the* Paw140 strain compared to that in the mt-2 strain (Fig. 2C).

### Exclusion of xyl mRNAs from the nucleoid is independent of the transcriptional machinery

A second type of perturbation entered in the architecture of the *xyl* gene expression flow involved complete replacement of the native *σ*^54^-dependent *Pu* promoter of the *upper* operon of the pWW0 plasmid (40, 41; Fig. 1A) by a heterologous expression T7 system. The replacement was entered in the pWW0 plasmid as explained in the Material and Methods section and the resulting construct (pTOL-PuxT7) then passed to strain *P. putida* KT2440•T7 (42), which expresses the T7 RNA polymerase through a genomic insertion of a *lacI*^*q*^*/P*_*lac*_*-T7pol* cassette (Fig. 6A). As shown in Supplementary Fig. S5A, the modified plasmid did not support *m*-xylene metabolism in cells without T7 RNAP because it did not transcribe the *upper* pathway. Due to the leakiness of the *P*_*lac*_*-T7pol* born by in the *P. putida* chromosome, some red signals indicative of x*ylUW* expression were detected as well in samples without IPTG (Supplemental Fig. S5C). Note also that there were RNA dots stemming from the *lower* pathway in the wild-type *P. putida* mt-2 host bearing the modified plasmid pTOL-PuxT7 (Supplementary Fig. S5B). This is because *m*-xylene-activated XylR caused overexpression of XylS, which in turn suffices to activate the *Pm* promoter even in the absence of any effector (5, 43, 44). As a consequence, when pTOL-PuxT7 was placed in *P. putida* KT2440•T7, [i] *P*_*T7*_ activation elicited formation of both *xylUW* transcripts (Fig. 6, panels 1-4) and *xylX* transcripts (Fig. 6, panels 5-8) in the complete absence of the aromatic inducers used before and [ii] cells could grow on minimal medium with *m*-xylene as sole carbon source (Supplemental Fig. S5A) due to the performance of both the *upper* and the lower TOL pathways. This made sense within the current model of regulation of the *xyl* operons discussed above (Fig. 1A). But in contrast to the effect of the native effectors in the naturally-occurring system, the surrogate control of the *upper* route by T7 polymerase originated a higher number of distinct foci per cell. While this reflected the strength of the *P*_*T7*_ promoter as compared to *Pu* (which propagated into a more potent activity of the *Pm* promoter as well), it is noteworthy that the red signals always appeared focused (i.e. constrained within a subcellular domain) rather that diffused through the cytoplasm. Furthermore, they again materialized mostly in the nucleoid free regions. Automated quantification of signals detected with the red and blue channels in individual cells with pixel precision revealed that 91% of *xylUW* and 98% of the *xylX* were excluded from the DAPI-stained field (Fig. 6C). Given that such transcripts originate in single promoters per bacterium and that nucleoid-free domains of *P. putida* cells are filled with ribosomes (20) it is possible that the images of Fig. 6 reflect the detachment of the transcripts from their promoters (whether *P*_*T7*_ for *xylUW* and *Pm* for *xylX*) and their resetting elsewhere in the cytoplasm for translation.

**FIG 6.**
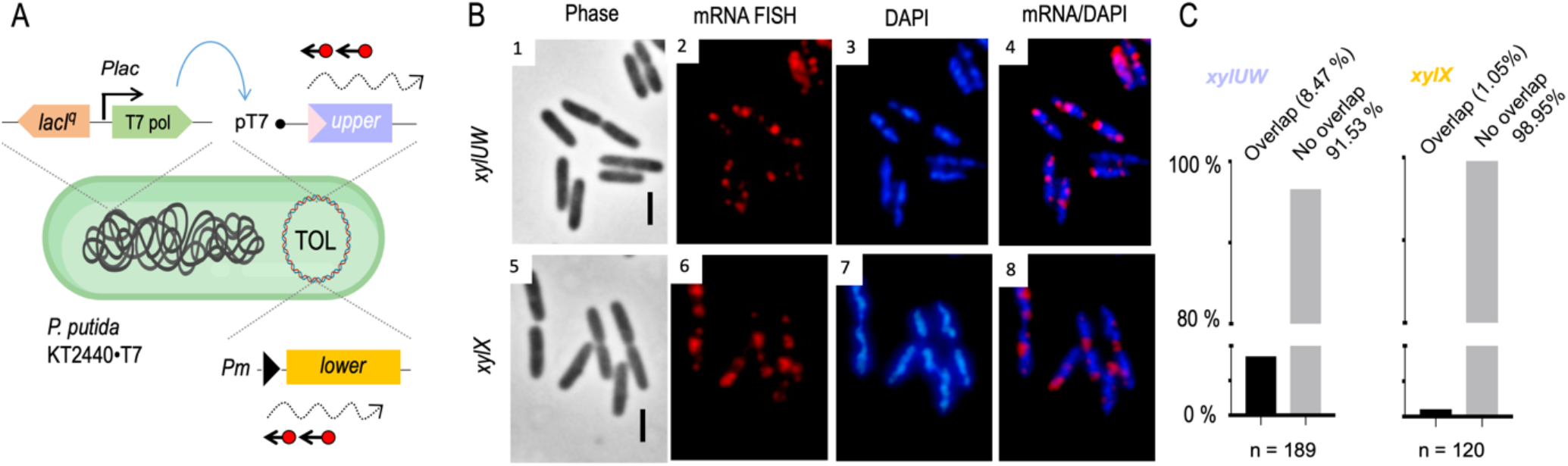
Localization of the *xyl* mRNA transcribed with an orthogonal RNAP. (A) Sketch of RNA-FISH experiments with the modified plasmid pTOL-PuxT7 in the KT2440•T7 strain, expressing T7 RNAP by the chromosomal *lacI*^*q*^*/Plac*-*T7pol* cassette. After exposure of the cells for 2 h with *m*-xylene, *xyl* mRNAs were visualized with the FISH. (B) Images obtained from the FISH experiment by using either the *xylUW* or the *xylX* probe set are shown: mRNAs (red; panel 2 and 6), DAPI-stained DNA (blue; 3 and 7) and merged signals are presented (panel 4 and 8) in the fixed cells (panel 1 and 5). Scale bar, 2.5 µm. (C) Quantification of overlapping regions between mRNA-red signals and DAPI-blue signals with sub-pixel precision. Over 100 exemplary cells were considered for the colocalization analysis.

In addition, one important evidence embodied in this set of experiments is that formation of mRNA foci and apparent motion towards ribosome-rich subcellular sectors occurs regardless of whether the cognate RNAP can (host polymerase) or cannot (T7pol) couple transcription to translation. This rules out that the distinct foci of *xylUW* and *xylX* mRNAs shown throughout this work reflect the action of transcription/translation complexes (the so-called expressome; 45, 46) on the nucleoid surface. Instead, it seems that the untranslated TOL transcripts move away from their promoters, plausibly towards other subcellular locations for translation. But, if that were the case, what physical forces prevent their diffusion through the cytoplasm?

### Disruption of intracellular crowding enables xyl transcripts to diffuse throughout the cytoplasm

The data above accredit that *xyl* mRNAs leave the proximity of the nucleoid towards the ribosome-enriched subcellular domains. Although a role for specific RNA binding proteins cannot be ruled out as drivers of the process, a simpler explanation is that the corresponding sequences endow the *xyl* transcripts with physical properties that make them to be quickly discharged from the vicinity of the nucleoid, especially if the transcribed sequences are large. In fact, it seems that smaller RNAs use to appear uniformly distributed throughout the bacterial cell, while longer molecules typically display more limited dispersion (47). Large mRNAs can hardly co-localize with densely packed DNA regions due to straight physical forces e.g. excluded volume effects (48, 49). As a consequence, the more crowded the bacterial cytoplasm is the less diffusible cellular components and molecules are, logically, in a size-dependent fashion (50, 51).

In order to test whether such a mutual exclusion between different intracellular domains accounted for the unusual behavior of the *xyl* mRNAs we treated *P. putida* KT2440•T7 (pTOL-PuxT7) cells with rifampicin. In one hand, since T7 RNAP is not sensitive to this antibiotic, the *upper* TOL transcript (*xylUW*) can still be produced (the *xylX* gene of the *lower* pathway predictably cannot, though). On the other hand, it is known that diffusion rate of ribosomal proteins is faster (11, 12) and the cDNA of the nucleoid expands after treatment with the drug (52, 53). Finally, the dearth of ribosomes available for interacting with RNAs due to inhibition of 16s rRNA synthesis with (54) may also ease diffusion of otherwise tethered transcripts. As a consequence, rifampicin treatment elicits a major change in the partition of the different components of the gene expression flow and causes a less compact intracellular milieu.

After treatment of growing *P. putida* KT2440•T7 (pTOL-PuxT7) for 2 h with the drug and *m-*xylene vapors, the mRNAs were visualized with FISH as before. Expectedly, we could hardly detect any *xylX* mRNA signals (Supplementary Fig. S6). In contrast, when cells were hybridized with the *xylUW* probe under the same conditions, red signals did appear (Fig. 7 and Supplementary Fig. S5). Yet, inspection of individual bacteria revealed a considerable variability in the intensity of the signals and their intracellular distribution. Some cells had their whole contour non-uniformly filled with red color, while other showed a number of discrete red signals with lower intensity (Fig. 7 and Supplementary Fig. S7). This outcome is not altogether unexpected, as the strong T7 promoter in the plasmid (55) drains cellular resources (56, 57) for the sake of transcription of the *upper* pathway, thereby originating noise. This effect can be exacerbated in a plasmid, as the loss of gyrase production with the drug and the ensuing accumulation of local supercoiling can lead to stochastic transcriptional bursts (58-60). In any case, we argue that the lack of distinct localization patterns in rifampicin-treated cells reflects the diffusion of the TOL transcripts under the conditions. Moreover, some images showed enrichment of *xylUW* mRNAs in the cells’ internal periphery in bacteria where the chromosome was otherwise highly compacted (Supplementary Fig. S7, marked). In these cases, it looked like mRNA was unable penetrate the denser DNA regions but could freely diffuse in the rest of the cytoplasmic space. Taken together, analysis of images shown in Fig. 7 and Supplementary Fig. S7 indicated that the *xylUW* transcript of rifampicin-treated cells lost its restraint in discrete foci and could then freely circulate as the cellular crowding decreases upon antibiotic addition. In sum, the data suggests that under native conditions *xyl* mRNAs become localized away from the nucleoid because of their entrapment with the translational machinery and the physical forces that determine phase separation between the different components of the gene expression flow. As shown above, if such a separation is perturbed upon rifampicin addition, *xyl* transcripts can then circulate through the whole cytoplasmic interior.

**FIG 7.**
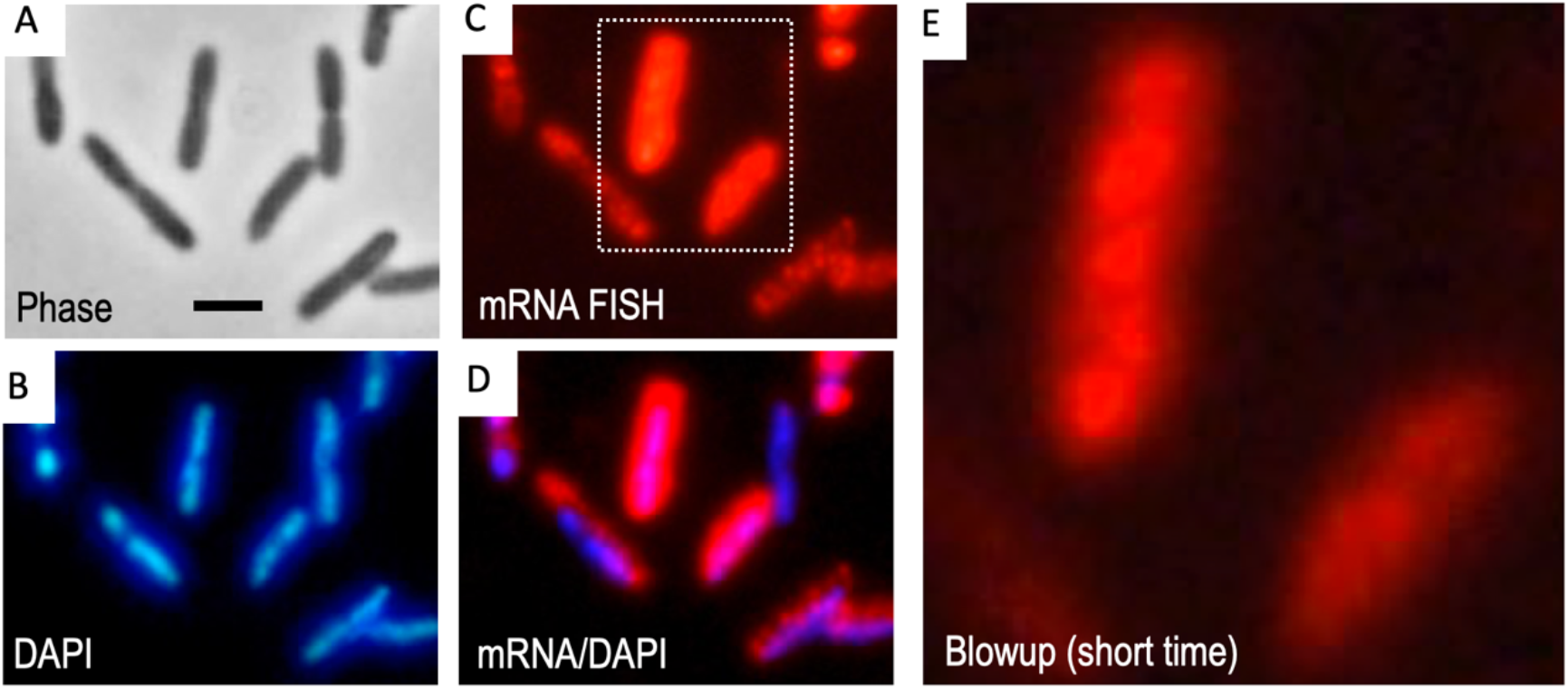
Effect of modulated macromolecular crowding by inhibiting bacterial transcription on the spatial distribution of the *xylUW* mRNA. The *P. putida* KT2440•T7 (pTOL-PuxT7) strain was tested to visualize the *xylUW* mRNA under transcription inhibited condition. After treatment of the cells for 2 h with *m*-xylene and rifampicin (200 µg ml^−1^), RNA-FISH experiment was proceeded. The resulting images were obtained as following: (A) Phase-contrast. (B) Red signals indicate fluorescently labeled oligos that hybridize to the *xylUW* mRNA, which are dispersed throughout the cell. (C) DAPI-staining. (D) The merged picture between mRNA-red and DAPI-blue channels. (E) The subset image of panel (C) was also taken with relatively shorter exposure time to rule out the blurred signals are not due to the light diffraction. Scale bar, 2.5 µm.

## Conclusion

This study shows that *xyl* mRNAs are constrained within subcellular regions with limited mobility rather than being freely diffusible inside *P. putida* mt-2 cells. Most *xyl* transcripts were detected in the space peripheral to the nucleoid, i.e. in a subcellular domain with virtually no DNA but enriched in ribosomes (20). That TOL mRNAs were detected away from the genetic loci where they originated suggests that they can migrate to translational sites from the 3D spot of the cell where they were synthesized. Our data show also that physical forces and translation is crucial for bringing about this scenario, which disappeared when in-house transcription was halted to reduce cellular crowding and ribosome engagement. This picture departs from the standard view of transcription-translation coupling, where RNAP and ribosomes get together in the so-called expressome complex (45, 46) which, obviously, must operate in close proximity to the nucleoid’s DNA encoding the gene(s) at stake.

While transcript entrapment is a known mechanism of RNA control in eukaryotes (61-63) the phenomenon as such has not been documented in prokaryotes. However, should transcript diffusion away from the promoter and capture by ribosomes in a different cell location occur, confinement in given spots for translation could become apparent. Some mRNAs can indeed travel throughout the *E. coli*’s interior (64) and translation inhibition makes RNA to remain closer to the nucleoid (47). This suggests that at least in some cases ribosomes in fact pull mRNA from the nucleoid towards the rest of the cytoplasm. Reality is that the interplay between the various components of the gene expression flow vary dramatically among bacterial types. In particular, the nucleocytoplasmic (NC) ratio changes from one greatly from one species to the other, what originates cytoplasms with different biophysical properties (65) that affect ribosome mobility and localization. For instance the high NC ratio of *Caulobacter crescentus* forces ribosomes to mRNAs and their translating ribosomes to remain close to their gene loci in the chromosomal DNA (11, 65, 66). In contrast the low NC ratio of *E. coli* allows segregation of ribosomes and mRNA away from their transcriptional sites. *Pseudomonas sp*. has also a low NC ratio (65) and the ribosomes are spatially segregated from the nucleoid (20). It is thus conceivable that TOL transcripts are not coupled to translation as they are produced but they instead move after complete transcription towards ribosome-rich cytoplasmic spots.

What could be the advantage of this setting of the *xyl* gene expression flow in *P. putida* as compared to an alternative subcellular architecture? This species has bona fide *nusA* and *nusG* homologues encoded in its genome. This indicates that transcription/translation coupling can indeed occur (45, 46), although its global incidence as compared to *E. coli* or *Bacillus* (19) is unknown. It may well happen that having movable *xyl* mRNAs eases assembly of the biodegradative complex encoded by the TOL operons for degrading aromatic compounds. This is because the cognate catabolic enzymes, in particular those which are membrane-bound (see above) could be synthesized at an optimal location site close to the site of action. Moreover, thereby synthesized enzymes could be produced close to each other and enable metabolic channeling (67). Finally, while Rho and transcription/translation coupling helps avoiding RNAP collisions and R-loops—with the downside of polarity—some bacterial types could have solved the traffic jam problem by evolving a different type of termination (19). In fact, interruption of non-translated transcripts may be detrimental for emergence of new catabolic operons: nosense mutations in lead genes could prevent expression of the whole cluster and curb evolution of downstream ORFs. These are all of course hypothetical scenarios that deserve further studies.

## MATERIALS AND METHODS

### Culture conditions

Unless otherwise indicated, *E. coli* and *P. putida* were routinely grown at 37°C and 30°C, respectively, in Luria–Bertani (LB) or M9 minimal medium (6 g l^−1^ Na_2_HPO_4_, 3 g l^−1^ KH_2_PO_4_, 1.4 g l^−1^ (NH_4_)_2_SO_4_, 0.5 g l^−1^ NaCl, 0.2 g l^−1^ MgSO_4_·7H_2_O) with 10 mM succinate. Bacteria were cultured in 100-ml Erlenmeyer flasks with shaking at 170 rpm with 20 ml of the medium specified for the corresponding experiment. Whenever necessary kanamycin (Km, 50 µg ml^−1^), ampicillin (Ap, 150 µg ml^−1^), gentamycin (Gm, 10 µg ml^−1^) or chloramphenicol (Cm, 30 µg ml^−1^) was added to cultures of bacterial cells for ensuring plasmid retention and maintenance of manipulated genotype. For the induction of TOL catabolic genes during RNA-FISH experiment, *P. putida* strains, carrying the *xyl* genes either in the pWW0 plasmid or in the chromosome, were overnight cultured in succinate amended M9 medium and then the bacterial culture were 100-fold diluted in the same medium and grown until exponential phase (OD_600_ = 0.3-0.5). Samples were then either cultivated further without additional substrate or exposed to vaporous *m*-xylene (1/2 dilution in dibutylphthalate, which is a non-effector for TOL genes) in a flask for 2h. To validate whether the output of the FISH approach matches the known TOL catabolic system, we cultured the mt-2 strain with other vaporous effectors such as toluene, *o*-xylene or soluble substrates such as benzoate (5 mM) and 3MBz (5 mM). The transcription of the *xyl* genes by bacterial RNA polymerase was halted by adding 200 µg ml^−1^ rifampicin (Rif) to the grown medium of the cognate sensitive strains.

### Construction of bacterial strains

The bacterial strains, plasmids, and primers used in this study are described in Supplementary Tables S1 and S2. General methods for DNA manipulation were followed standard protocols (68). Plasmid DNA was isolated from bacterial cells using commercial Wizard *Plus* SV Minipreps DNA Purification kit (Promega) or the QIAprep Spin Miniprep Kit (Qiagen). To replace the *Pu* promoter of the *upper* operon of the pWW0 plasmid by the orthogonal transcription T7 system, we used the seamless allelic replacement method of (69). The delivery plasmid for executing the promoter replacement pTOL-*Pu*x*T7* was built as follows. First, the upstream (TS1^*Pu*^, ∼0.5 kb) and downstream (TS2 ^*Pu*^, ∼0.5 kb) regions of the pWW0 plasmid around the *Pu* promoter were amplified with primer pairs PuxT7-TS1F/R and PuXT7-TS2F/R respectively (Supplementary Table S2). SOEing PCR then joined the amplicon TS1^Pu^ and TS2^Pu^, including 3’- and 5’ theT7 promoter sequence overhang complementary, by using primer pairs PuXT7-TS1F and PuXT7-TS2R (Supplementary Table S2). The resulting PCR product was cloned into the pEMG by using restriction enzymes such as *Eco*RI and *Bam*HI and ligation within the same site of the vector yielded the plasmid pEMG-PuxT7, which was kept into the *E. coli* DH5*α λpir* strain. This plasmid was transferred to *P. putida* mt-2 by triparental mating using the *E. coli* HB101 (pRK600) as helper strain (70, 71). Next, the pSW plasmid that expresses I-SceI endonuclease under the *Pm* promoter (72) was introduced by electroporation into the pEMG-PuxT7 cointegrated strain and thus had resistance for both Km and Ap. The clones were grown in LB medium with Ap (500 µg ml^−1^) and 3MBz (15 mM) to activate the *Pm* promoter, allowing I-SceI expression. The cells were plated on LB agar and we confirmed the promoter replacement in the TOL plasmid, by testing the loss of the pEMG-PuxT7, encoded Km resistance gene. Km-sensitive clones were selected and PCR further confirmed the emergent colonies with T7F/ PuXT7-TS2R primer pairs (Supplementary Table S2).Then, the manipulated plasmid pTOL-*Pu*x*T7* was either maintained in the mt-2 strain or transferred into the KT2440•T7 strain by conjugation.

To label the pWW0 plasmid with the tandemly repeated *tet* operators, we inserted the arrays into the *orf105* locus of the plasmid. To this end, the 570-bp of PCR product was obtained using primer pairs 105F/ 105R (Supplementary Table S2) corresponding to the 5’ portion of the *orf105* gene of the pWW0 plasmid. Then, the amplified fragment was cloned into the *Hind*III and *Not*I cloning site of the pP30D-FRT-tetO vector (26), generating pP30D-FRT-*tetO*-orf105. The construct was either kept into the CC118 strain and subsequently transferred into *P. putida* mt-2 strain, in turn the backbone of the vector, including the *tetO* arrays, was integrated into the pWW0 plasmid, generating the strain mt-2 (pTOL-*tetO*).

### Fluorescent *in situ* hybridization

To visualize *xyl* mRNAs, RNA-FISH was performed as described previously (11) with some modifications. To this end, cells growing in culture medium with or without effector for *xyl* genes were fixed in formaldehyde solution (4% formaldehyde and 30 mM NaHPO_3_ pH 7.5) by incubating for 15 min at room temperature, followed by 30 min further on ice. Samples were collected by using centrifuge for 3 min at 4,500 g and removed supernatant. The cell pellets were washed with diethyl pyrocarbonate (DEPC)-treated phosphate-buffered saline (PBS) and resuspended the cells in 100 µl of GTE buffer (50 mM glucose, 20 mM Tris-HCl pH 7.5, 10 mM EDTA pH 8). Then, 12 µl of the mixture was transferred to 4 µl of the lysozyme solution containing lysozyme and vanadyl ribonucleoside complex (VRC), resulting in final concentration as 2.5 µg ml^−1^ of lysozyme and 2 mM VRC respectively. Next, 3 µl of the mixed solution was applied onto poly-L-lysine coated cover slip, and stored at room temperature to be dried on the cover slip. After putting the cover slip into a rack, this assembly was immersed sequentially in methanol for 10 min and acetone for 30s at −20 °C. Once the coverslip was dry, it was kept at 37 °C in 10% formamide solution (10% formamide, 2X saline-sodium citrate buffer (SSC) in DEPC treated water, 2 mM VRC). After 60 min, the solution was removed and 50 µl of the hybridization solution (10 % formamide, DEPC treated 2X SSC, 10 % dextransulphate, 2 mM VRC, 40 U RNase inhibitor, and 250 nM CAL Fluor Red 610 labeled FISH probes; Stellaris™, Biosearch Technologies, Supplementary Tables S3) was spotted onto the cover slip and the sample was incubated in a dark and humid chamber at 42 °C for the hybridization process. After overnight, the sample was then washed twice with the solution (10% formamide and DEPC treated 2X SSC) for 15 min at 37 °C and DAPI (2.5 ug ml^−1^) staining was performed in second washing step. After brief rinse with PBS, the coverslip was assembled with slide glass including antifade reagent Prolong (Invitrogen) and sealed by clear nail polish. The specimen was visualized by fluorescent microscope. To identify cellular position of the TOL plasmid carrying the tandem copies of the *tet* operators, DNA-FISH was exploited with the 6-carboxyfluorescein (FAM) labeled locked nucleic acid (LNA) *tetO* probe (5’ 6-FAM-CTCTATCACTGATAGGGA; Bionova). It is the same strategy as RNA-FISH, but the hybridization was carried out at two different temperatures: at 95 °C to denature DNA for 2 min and followed by at 42 °C overnight in a dark and humid chamber. The fluorescent signals representing *xyl* mRNAs and the *tetO*-tagged DNA were achieved by employing sequentially combined RNA/ DNA-FISH. We followed the same procedure the RNA-FISH with *xyl* probe sets prior to performing the DNA-FISH with the *tetO* probe.

### Microscopy and image analysis

Microscopy was performed using an Olympus BX61 apparatus equipped with *×*100 phase contrast objective and a DP70 camera of the same brand. Signals for red-RNA, DAPI, and green-DNA were obtained using wide field excitation with following filters; MWIY2, U-MNU2, and U-MNIBA2. Process of multi-channel images obtained from the microscopy and measurement of relative fluorescent intensities were carried out by using the software Fiji. To analyze fluorescent foci representing mRNA molecules in terms of their relative cellular positions in respect of the DAPI stained-nucleoid, we used the Cellshape (23). To analyze overlap between the red and the blue signals, the contour lines of both channels were calculated over the RGB TIF formatted images. Firstly, pixel values (red/blue channels) were extracted. Secondly, the resulting values were normalised within the interval [0,1] in order to allow for direct channel comparison regardless any potential technical inconsistencies in measuring any one particular signal. Finally, contour lines were calculated with sub-pixel accuracy. The overlap value refers to the overlap between the contours for the red signal and the contours for the blue signal

## SUPPLEMENTAL MATERIAL

*Supplemental material for this article may be found on line at* XXX

**Supplementary Tables** S1, S3 and S3

**Supplementary Figures** S1, S2, S3, S4, S5, S6 and S7

## ACKNOWLEDGEMENTS

V.D.L. and J.K. designed this study and the experimental layout. J.K. did all the experiments with the assistance of A.G-M, who developed and implemented the software for image analysis used throughout this work. J.K. and V.D.L. wrote the manuscript with further contributions from A.G-M. to data analysis and interpretation of the results. We thank Christine Jacobs-Wagner for sharing the protocol for FISH approach. Isabelle Vallet-Gely and Karl-Erich Jaeger are gratefully acknowledged also for generous sharing of valuable materials, as is Belén Calles for insightful discussions. This work was funded by the SETH (RTI2018-095584-B-C42) (MINECO/FEDER) and SyCoLiM (ERA-COBIOTECH 2018 - PCI2019-111859-2) Projects of the Spanish Ministry of Science and Innovation, the MADONNA (H2020-FET-OPEN-RIA-2017-1-766975), BioRoboost (H2020-NMBP-BIO-CSA-2018-820699), SynBio4Flav (H2020-NMBP-TR-IND/H2020-NMBP-BIO-2018-814650) and MIX-UP (MIX-UP H2020-BIO-CN-2019-870294) Contracts of the European Union and the InGEMICS-CM (S2017/BMD-3691) Project of the Comunidad de Madrid - European Structural and Investment Funds - (FSE, FECER). Authors declare no conflict of interest.

