## Supplementary Figures for "The subcellular architecture of the *xyl* gene expression flow of the TOL catabolic plasmid of *Pseudomonas putida* mt-2"

**Supplementary FIG S1.** RNA-FISH experiments with *xyl* probe sets in the mt-2 strain grown with/ without TOL aromatic effectors

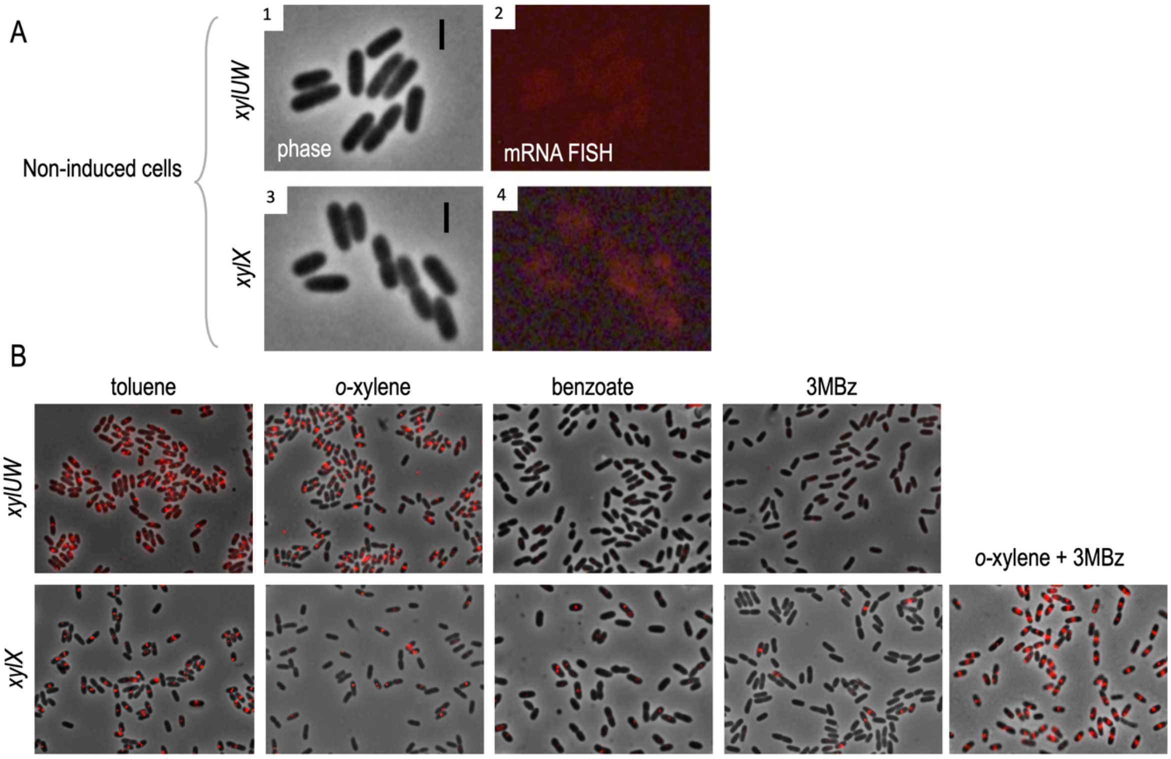


(A) RNA-FISH microscopy with non-induced cells. No detectable RNA signals were observed in the red channel from the approach. Phase-contrast images (panel 1 and 3) and their counterpart red channels representing *xyl* mRNAs (panel 2 and 4) are shown. Scale bar, 2.5 µm. (B) Stacked phase contrast and FISH cross-section images of the cells, cultivated with toluene, o-xylene, benzoate (5 mM), or 3MB (5 mM) for 2 h. Red fluorescently labeled oligos that hybridize to either the *xylUW* or the xylX mRNA were used in the experiment. The culture condition with mixed effectors (*o*-xylene + 3MBz) was also tested to enhance the expression of the *lower* pathway.

**Supplementary FIG S2.** DNA-FISH to visualize the pWW0 plasmid tagged with tandem copies of *tet* operators

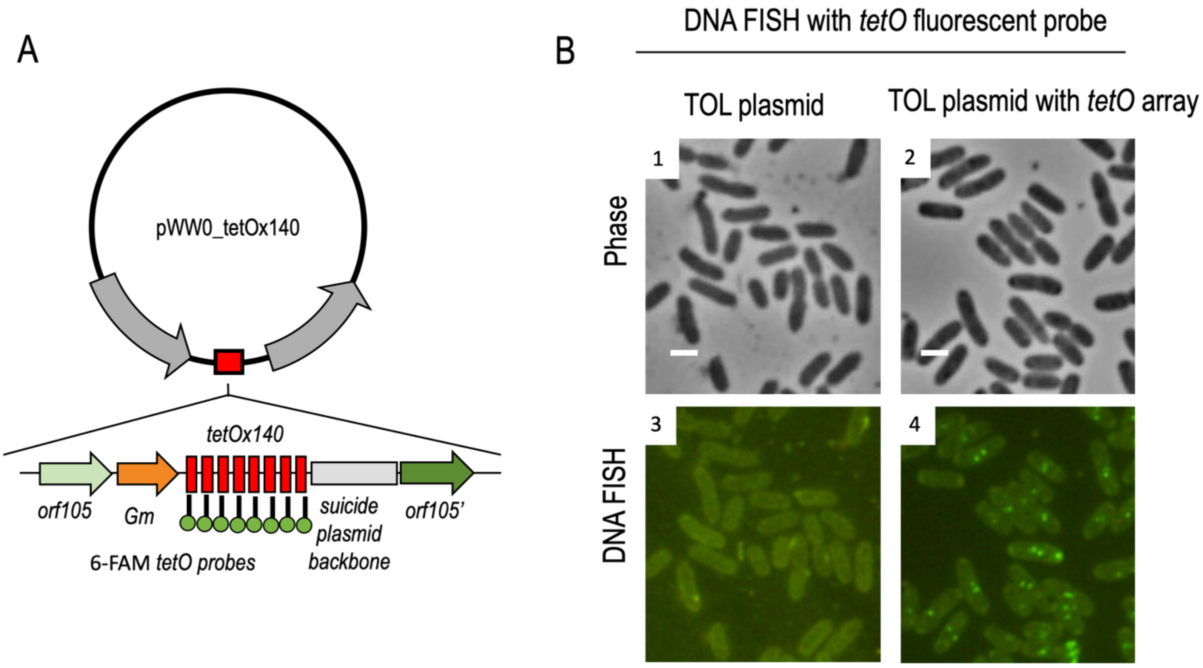


(A) Schematic representation of the modified pWW0 plasmid carrying tandemly repeated *tetO* arrays in the *orf105* locus. The 6-Carboxyfluorescein (6-FAM)-labeled *tetO* probe (LNA structure) was used in DNA-FISH approach to sense the plasmid DNA. (B) The hybridization was proceeded with the probe on fixed cells carrying either the intact TOL plasmid or the *tetO* array labeled pWW0 plasmid. As a result of the FISH microscopy, phase-contrast (panel 1 and 3) and DNA-green signals (panel 2 and 4) were obtained. Scale bar, 1 μm.

**Supplementary FIG S3.** Dual labeling of the pWW0 plasmid and *xyl* mRNAs

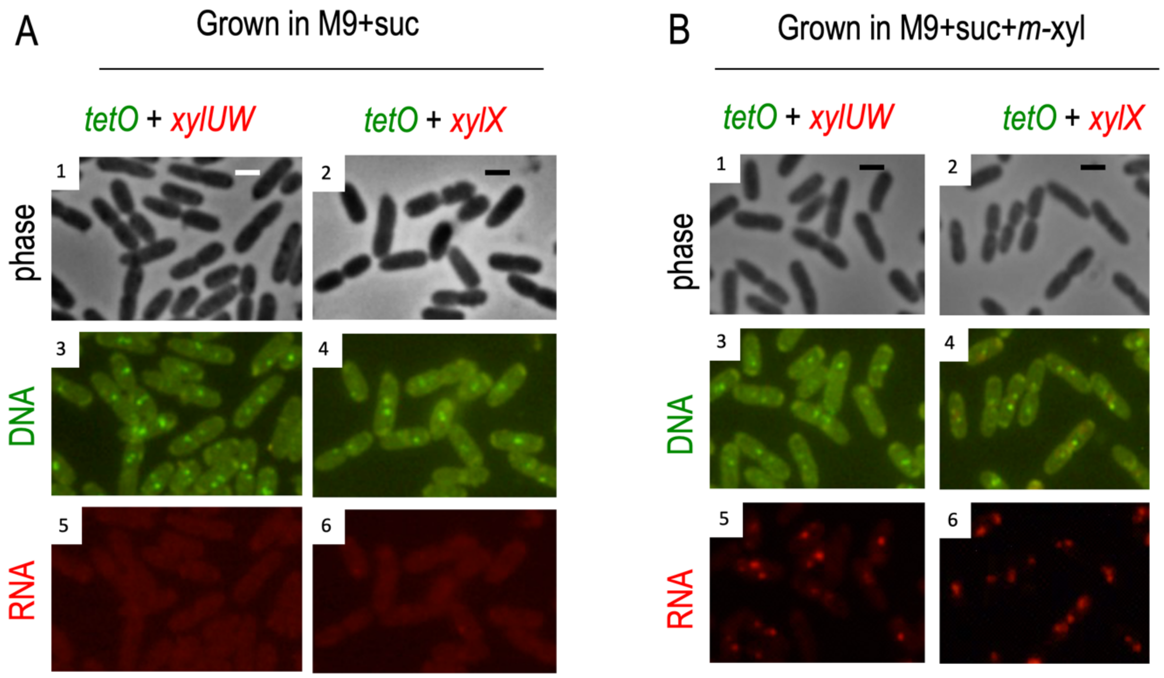


(A) Using the mt-2 (pTOL-t*etO*) strain, sequentially combined RNA-DNA FISH was conducted with fixed cells grown without effector. (B) The same approach was applied to the cells exposed to the aromatic effector *m*-xylene. The combined FISH experiment enabled to detect the plasmid DNA (green signals; panel 3 and 4) and *xyl* mRNAs (red signals; panel 5 and 6) in the cells (panel 1 and 2) dependent upon induction of the TOL catabolic system. Scale bar, 1 µm.

**Supplementary FIG S4.** Single-cell mapping of the *xyl* gene expression flow

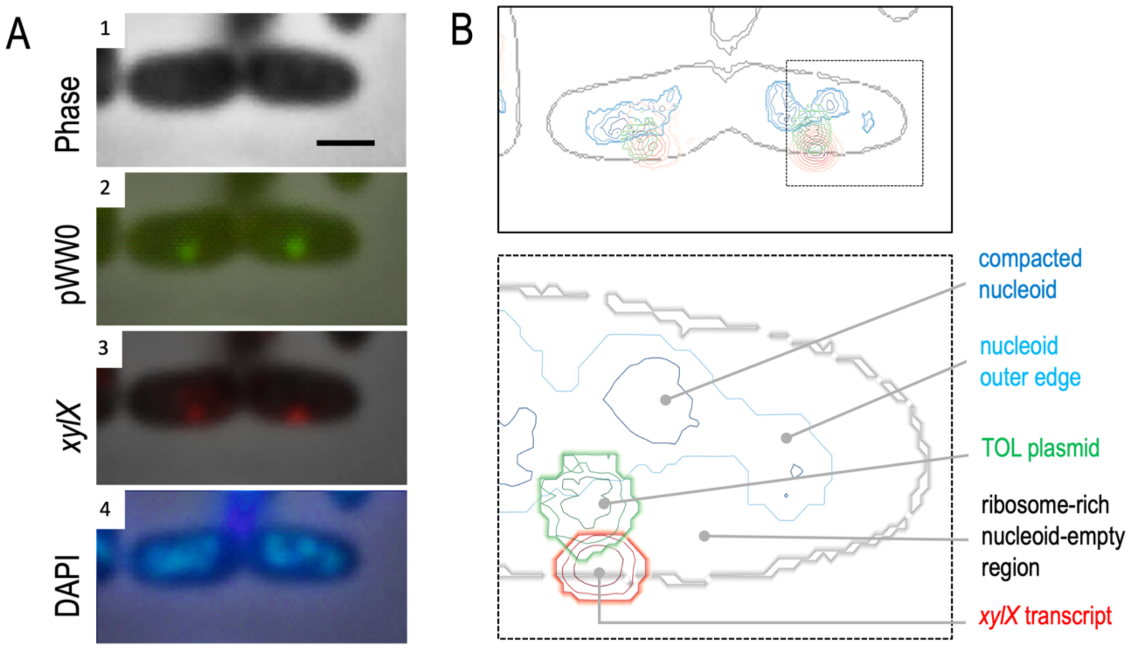


(A) An exemplary cell, from the combined RNA-DNA FISH microscopy, shows signals representing the pWW0 plasmid (green; panel 2), the *xylX* mRNA (red; panel 3), and the nucleoid (blue; panel 4), respectively. Each fluorescent channel was overlaid on the phase-contrast image (panel 1). Scale bar, 1 µm. (B) All the channels, which appeared in panel A, were merged using the image analysis tool (upper panel) and the blow-up picture (lower panel) demonstrated the subcellular localization of each molecule. While the plasmid DNA is linked to the nucleoid, the *xylX* transcript is found in the peripheral space of the cytoplasm.

**Supplementary FIG S5.** Expression of the *xylUW* mRNA with the orthogonal expression T7 system and visualization of the *upper* transcripts

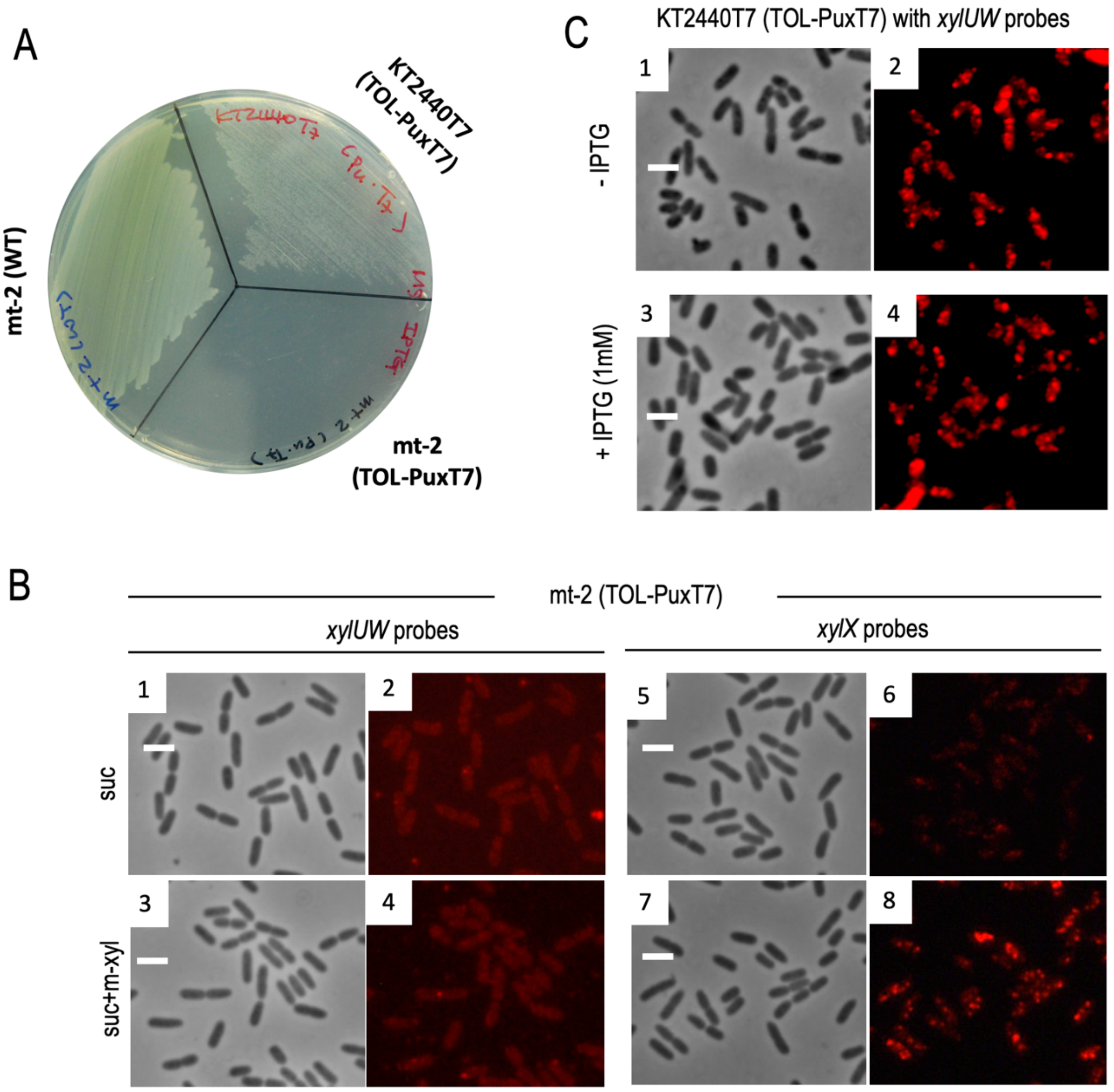


(A) Cells were grown on M9 minimal solid medium containing 1 mM IPTG with vaporous *m*-xylene as a sole carbon source; strains such as mt-2 (WT), mt-2 (pTOL-PuxT7), and KT2440•T7 (pTOL-PuxT7) were examined. (B) RNA-FISH was carried out with the modified plasmid pTOL-PuxT7 in the strain, where do not carry the T7 RNA polymerase. The resulting images were obtained from the experiment with the cells grown in succinate-supplemented minimal medium (panel 1, 2, 5 and 6). Also, the cells exposed to *m*-xylene were examined with the same approach. The phase-contrast images (panel 3 and 7) and their counterpart red channels displaying *xyl* mRNAs (panel 4 and 8) are shown. (C) Labeling of the *xylUW* mRNA in the KT2440•T7 (pTOL-PuxT7) strain. Red outputs, representing the *xylUW* mRNA, were detected irrespective of IPTG treatment (panel 2 and 4), enhancing expression of the T7 RNA polymerase. Scale bar, 2.5 μm.

**Supplementary FIG S6.** Visualization of the *xylX* mRNA under inhibition of bacterial RNAP

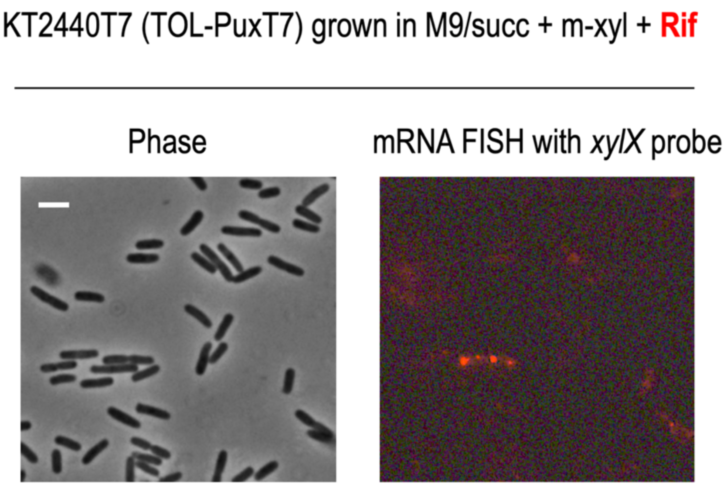


After treating the culture of the KT2440•T7 (pTOL-PuxT7) strain for 2 h with *m*-xylene and rifampicin (200 µg $ml^{-1}$), the cells were processed for the FISH experiment with the *xylX* probe set. Expectedly, few RNA-red signals were detected due to the inhibition of bacterial RNAP. Scale bar, 2.5 µm.

**Supplementary FIG S7.** The FISH microscopy images demonstrated in Figure 7 with more examples.

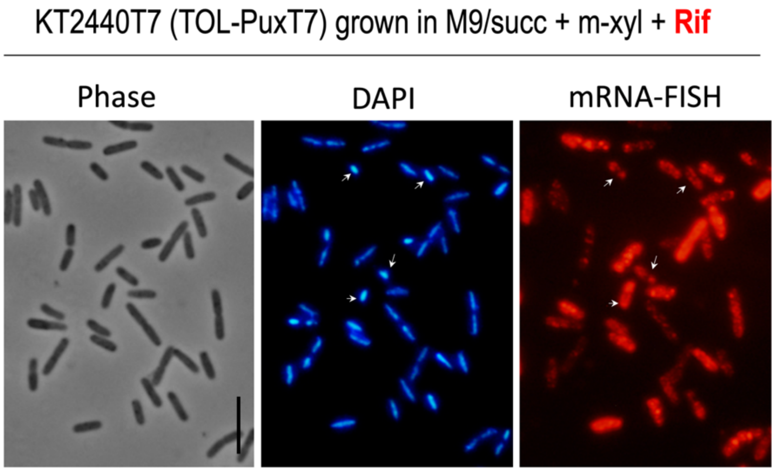


The fluorescent intensity of RNA-red outputs against the *xylUW* transcripts are varied from cell to cell when rifampicin treated to the cells; some cells show very high signals, whereas the other cells display relatively low or middle intensities. Also, when the nucleoid was compacted by the drug, the mRNA majorly accumulated in the peripheral space of the cytoplasm (marked with arrows). Scale bar, 5 μm.
