## Supplementary Tables for "The subcellular architecture of the *xyl* gene expression flow of the TOL catabolic plasmid of *Pseudomonas putida* mt-2"

**Supplementary Table S1.** Description of bacterial strains and plasmids used in this study

| Strain | Description | Reference |
|---|---|---|
| *P. putida* | | |
| mt-2 | *P. putida* wild type harbouring the pWW0 plasmid | (1) |
| PaW140 | *P. putida* carrying the *upper* and the *lower* operons in the chromosome with corresponding regulator genes such as the *xylR* and the *xylS* genes | (2) |
| KT2440•T7 | $Gm^R$, KT2440 derivate with chromosomal T7 polymerase expression system | (3) |
| mt-2 (pTOL-*PuxT7*) | mt-2 strain carrying the T7 promoter for the *upper* pathway in the pWW0 plasmid | This study |
| KT2440•T7 (pTOL-*PuxT7*) | KT2440•T7 strain carrying the T7 promoter for the *upper* pathway in the TOL plasmid | This study |
| mt-2 (pTOL-tetO) | mt-2 strain carrying the pWW0 plasmid tagged with tandem copies of the *tet* operators | This study |
| | *E. coli* | |
| HB101 | $Sm^R$, *hsdR-M*$^+$, *pro, leu, thi, recA* | (4) |
| DH5α*λ pir* | *λ pir* phage lysogen of DH5α | Lab collection |
| CC118 | *F-, Δ(ara-leu)7697, araD139, Δ(lac)X74, phoAΔ20, galE, galK, thi, rpsE, rpoB* | (5) |

| Plasmids | Description | Reference |
|---|---|---|
| pRK600 | $Cm^R$, *oriV* ColE1, *tra*$^+$*mob*$^+$ of RK2, helper plasmid for mobilization in tripartite conjugations | (6) |
| pSW | $Ap^R$, *ori*RK2, *xylS,* bearing a *Pm* → *I-sceI* transcriptional fusion | (7) |
| pEMG | $Km^R$, *ori*R6K, suicide plasmid with two I-SceI sites flanking the *lacZ*α polylinker | (8) |
| pEMG- PuxT7 | $Km^R$, pEMG carrying T7 promtoer between the upstream and downstream flanking regions of the Pu promoter in pWW0 plasmid | This study |
| pP30D-FRT-tetO | $Gm^R$, ColE1, template for cointegration of t*etO* arrays | (9) |
| ppD30FRP-teto-orf105 | $Gm^R$, pPD30D-FRT-tetO carrying the *orf105* gene between *Hind*III and *Not*I sites of the plasmid | This study |

**Supplementary Table S2.** Oligonucleotide primers used in this study

| Name | Sequences (5'-3') |
|---|---|
| PuxT7-TS1F | CGCGAATTCGTCGGATACGGCGGGCGACCG |
| PuxT7-TS1R | CCCTATAGTGAGTCGTATTAAGAAGACAGCCTTG ACTTTCA |
| PuxT7-TS2F | TTAATACGACTCACTATAGGGGACTTAAAATAAA AATAGTG |
| PuxT7-TS2R | CGCGGATCCCAAATGTTATAGGTAGCAAGGA |
| T7F | TAATACGACTCACTATAGGG |
| 105 F | CGCAAGCTTATGAGCGATCCGGCCGTCG |
| 105 R | CGCGGTACC GCGTCCCGCGCCGAAGCGGC |

**Supplementary Table S3.** Oligonucleotide probes for RNA-FISH

| *xylUW* probe set* | *xylX* probe set* |
|---|---|
| AGCAACCAATCTGAACAGAG | AGGTGCATTGTCATGGTCAT |
| CCCGCTTTGAGGATATACAT | ACTATCTATATAGTCGAGCC |
| TCACAGACTCCAGGCGTAAC | TAGATGCCCTCGTTCTCATC |
| CTCAGAAAGCACTAGGCCAG | GAACATCTCGCGCTTGCAGC |
| CTCACCAAAATTGGTGGTCG | AATCGAACAGCCGAGGGTCG |
| ATAACTGCGACGAAAATGGT | TCAAAGATGTGTTTCATCTC |
| CCTATCACGAGAGATGAAGC | GGCGAGATAAATCCAGTTGC |
| CGTTGGACTGGCATCTATAA | GTTCTTCTCGGGAATCTGGC |
| ATTGAAGATTGATGCAGCCG | CCATCTGCGTGGTGTAATAG |
| CTGGGCATATAGTCGGTTGA | TGTGATGAATATCGGCTGCC |
| CAGGCTGGATATATCGTTGC | TCAGCTCACCATCTTTGTTG |
| GGTAGATGACTAAGGCTCGA | GACTGCAGGCATTGACGAAG |
| TAGTAATGTCGCTGCAGCTG | CACTCCTAAAGCGACAGAGC |
| TTCCGAGATCGACACGACTA | CCGAATTGCTGAAGGTCCAG |
| AAGTTCTCGGCAACAACACG | TCTTTGACCTTGAGCAGCTT |
| ACTTAATGCATCACATGCAG | AGTCGAAGCTGTCCGGATAG |
| TTATAGGTAGCAAGGACGGC | TTCTTCAGGTCGTGCGAGCC |
| GCGAGCATTGAATCACCTAT | GTAGGAAGCAAAGCGCGCAA |
| GCCCTGCTTTAGTTTTCCTT | CAGGCTGCCGAATAGAAATC |
| CACATCATCGACAGATAGCC | GACTCGCCGAGGAACTCTTC |
| AGTACTGTGGGCCTCTTTAG | GACCATGTCGATGACCTTCC |
| AGTCAGTACCACAGATACCG | AGCACTTCCAGACCTTCGGG |
| TCATTCTTTTGGCGAACTCG | AAACATAGGTACTGGAACCG |
| CGCTAACTTCATGACCCAAG | TGCACTTTCCAGTTGCCTTC |
| GATCCGACCTGTTCAATCAC | TACTGACGTGGTAGCCGTCG |
| CAGGCCTTAGACCTTTAACG | GCGGCGTAGTTCCAGTGAAC |
| AGATGACTCTCCAGACTGAC | TCTCTCAGCTTGCGCTGCTG |
| CGTGTAGCACGTTCCACAAG | GTCATGGCGCGAATATCATC |
| CGTCTTAGGACAAACATGCG | TTCAAAGGAGTAGAAACCGC |
| GAAGCCTCCATCAAAGTCGA | GTGCCCAGACCATCTGGTGG |
| TTTCCGGAACCACAACGAAA | GGCGGTTTTTCGGGTCACCC |
| CGCAACCAGAAACAAGAACA | CGATCTCGCTCGGCGAACAG |
| TGCAATGTTCATAAGTCCGA | TTCACCAAACTCGCTGGCTA |
| GATGAATGTCCGGTGCAATG | GAGACGCCGATCATCCAGTC |
| ATTTTGGCTGCTGCAGTGAG | GAACTGGTCCATCAGGTAGA |
| GCATTAATGCATTCATCCGC | GACGGGTGATACGCAACTGC |

| | |
|---|---|
| CGCACAGTCTTATAAACGCT | GATTTCGGTTCTATCCACCG |
| GCTTGCCCAGAATAGTCAAT | CTCGGCGTTTCGCCTTTGGG |
| AATAAGCTCCTTTAACGCCG | GTCCTCGTACTGACGGACAC |
| AAATATCATGTGACGGGACG | CCATGCCGCTGACATTGAAG |
| CCACGTACAATAAGGCCTTT | GGAATTCCTCCAGGTCGTCC |
| ATCTTTCATACAGACGCCTG | CACGGGACATGTCGTTCATC |
| CCAATAAGCTAGTTGAACGC | CCCTCGATCCAGTGTTTGGC |

*Each oligo is labeled with the fluorophore CAL Fluor Red 610
